# TAPAS: A Thresholding Approach for Probability Map Automatic Segmentation in Multiple Sclerosis

**DOI:** 10.1101/609156

**Authors:** Alessandra M. Valcarcel, John Muschelli, Dzung L. Pham, Melissa Lynne Martin, Paul Yushkevich, Peter A. Calabresi, Rohit Bakshi, Russell T. Shinohara

## Abstract

Total brain white matter lesion (WML) volume is the most widely established magnetic resonance imaging (MRI) outcome measure in studies of multiple sclerosis (MS). To estimate WML volume, there are a number of automatic segmentation methods, yet, manual delineation remains the gold standard approach. These approaches often yield a probability map to which a threshold is applied to create lesion segmentation masks. Unfortunately, few approaches systematically determine the threshold employed; many methods use a manually selected threshold, thus introducing human error and bias into the automated procedure. In this study, we propose and validate an automatic thresholding algorithm, Thresholding Approach for Probability Map Automatic Segmentation in Multiple Sclerosis (TAPAS), to obtain subject-specific threshold estimates for probability map automatic segmentation of T2-weighted (T2) hyperintense WMLs. Using multimodal MRI, the proposed method applies an automatic segmentation algorithm to obtain probability maps. We obtain the true subject-specific threshold that maximizes Sørensen-Dice Similarity Coefficient (DSC). Then the subject-specific thresholds are modeled on a naive estimate of volume using a general additive model. Applying this model, we predict a subject-specific threshold in data not used for training. We ran a Monte Carlo-resampled split-sample cross-validation (100 validation sets) using two data sets: the first obtained from the Johns Hopkins Hospital (JHH) on a Philips 3 Tesla (3T) scanner (n = 94) and a second collected at the Brigham and Women’s Hospital (BWH) using a Siemens 3T scanner (n = 40). By means of the proposed automated technique, in the JHH data, we found an average reduction in subject-level absolute error of 0.1 mL per one mL increase in manual volume. Using Bland-Altman analysis, we found that volumetric bias associated with group-level thresholding is mitigated when applying TAPAS. The BWH data showed similar absolute error estimates using group-level thresholding or TAPAS likely since Bland-Altman analyses indicate no systematic biases associated with group or TAPAS volume estimates. The current study presents the first validated fully automated method for subject-specific threshold prediction to segment brain lesions.

## 1. Introduction

Multiple sclerosis (MS) is a chronic inflammatory and degenerative disease of the central nervous system characterized by demyelinating lesions occurring in the brain and spinal cord (Confavreux and Vukusic 2008; Compston and Coles 2002). The disease is associated with multifocal lesions and atrophy in brain white and gray matter leading to physical disability, cognitive dysfunction, and even unemployment (Rovira and León 2008; Tauhid et al. 2015). In MS research, diagnosis, and therapeutic monitoring, magnetic resonance imaging (MRI) is a commonly used tool to detect disease activity and quantify disease severity (Ge 2006; Zivadinov and Bakshi 2004; Bakshi et al. 2005). MRI allows for the detection of T2-weighted (T2) hyperintense white matter lesions which can be used to calculate and track important MS metrics such as lesion volume and count (Ge 2006; Dworkin et al. 2018). Typically, total lesion burden, or lesion load, is defined as the volume of total brain matter containing lesions, and is a cornerstone for assessing disease severity in MS research and clinical investigations (Popescu et al. 2013; Calabresi et al. 2014; Tauhid et al. 2014).

To quantify lesion burden, different approaches use MRI to identify and segment lesional tissue. Manual segmentation is the gold standard approach and requires an imaging expert or neuroradiologist to inspect scans visually and delineate lesions. Due to difficulties associated with manual segmentation such as cost, time, and large intra- and inter-rater variability, many automatic segmentation methods have been developed (Egger et al. 2017; Carass, Roy, Jog, Cuzzocreo, Magrath, Gherman, Button, Nguyen, Prados, et al. 2017; García-Lorenzo et al. 2013; Lladó et al. 2012). Unfortunately, since lesions present heterogeneously on MRI scans, automatic segmentation remains a difficult task, though numerous methods have been proposed. No single approach is widely accepted or proven to perform optimally across lesion types, scanning platforms, and centers. A common key step in automatically delineating lesions and thus measuring lesion volume involves creating a continuous map indicating the degree of lesion likelihood using various imaging modalities (Sweeney et al. 2014, 2013; A. M. Valcarcel, Linn, Vandekar, et al. 2018; Roy et al. 2015). In these cases, a threshold is then applied to probability maps to obtain binary lesion segmentations, also referred to in the field as lesion masks.

It has been reported anecdotally that automatic approaches may be susceptible to biases in lesion volume estimation associated with the total lesion load; that is, in subjects with few lesions automated techniques tend to over-segment lesions, and in subjects with higher lesion load, lesions are under-segmented. To investigate this, we leveraged the 2015 Longitudinal Lesion Challenge (https://smart-stats-tools.org/lesion-challenge) (Carass, Roy, Jog, Cuzzocreo, Magrath, Gherman, Button, Nguyen, Bazin, et al. 2017; Carass, Roy, Jog, Cuzzocreo, Magrath, Gherman, Button, Nguyen, Prados, et al. 2017), a publicly available data set consisting of five subjects for training and fourteen unreleased subjects for testing. In training and testing sets, subjects had at least four imaging visits. The training data contains manual delineations from two expert raters while the testing set does not publicly provide manual delineations; rather, the testing set only consists of volume estimates from each rater. Challengers who wish to compare new segmentation methods can submit their testing set automatic segmentations. The automatic segmentation method is ranked using a weighted average of various similarity measures. A leader board with method performance measures is maintained by challenge organizers and some published work compares top performing methods (Carass, Roy, Jog, Cuzzocreo, Magrath, Gherman, Button, Nguyen, Prados, et al. 2017).

We present data from challengers as Bland-Altman plots (Bland and Altman 2007, 2016) to assess disagreement with manual volumes from the top two performing approaches described in Carass, Roy, Jog, Cuzzocreo, Magrath, Gherman, Button, Nguyen, Prados, et al. (2017) (see appendix Table C3). Bland-Altman plots are provided in **Figure 1** to compare the automatically generated and manually delineated volumetric measures. This graphical approach presents the differences between techniques, automatic and manual, against the averages of the two. Horizontal lines are drawn at the mean difference and at the mean difference plus and minus 1.96 times the standard deviation of the differences, which are defined as the limits of agreement. Points found outside the limits of agreement indicate the difference between techniques is not clinically important and the two methods can be used interchangeably.

**Figure 1:**
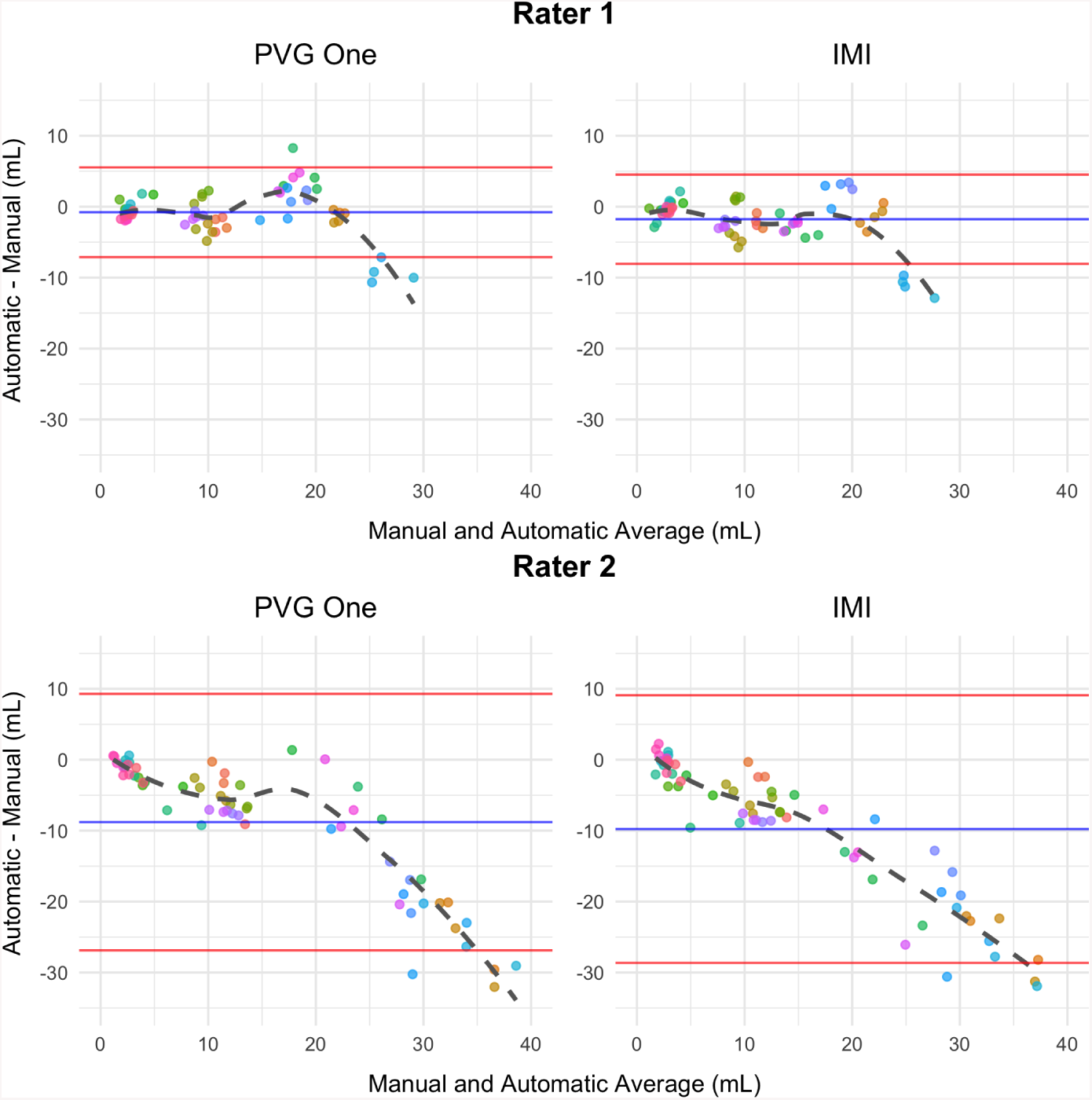
Bland-Altman plots using the first (left) and second (right) ranked automatic segmentation methods’ volumes from the 2015 Longitudinal Lesion Challenge are presented. We summarize volumes obtained from both rater 1 (top) and rater 2 (bottom). Using the differences, we highlight the mean (blue) plus and minus 1.96 times the standard deviation (red). Each subject is represented in a unique color and each point represents a subject-time point. There are five unique subjects with at least four follow-up imaging sessions.

The plots in **Figure 1** show systematic deviations in automatic and manual volumes. Both ranked methods show that, as lesion load increases, automatic segmentation approaches underestimate volume compared with rater 1 and rater 2. This is evident by the dashed fitted smooth line deviating away from the mean and outside the limits of agreement starting around lesion loads larger than 20 mL apparent in all four of the plots. While the direction of over- or under-estimation and magnitude varied for rater 1 and rater 2 across challenge submissions, each approach shows systematic deviation and bias in volume estimates.

The bias present in the volumetric estimates may be related to the thresholding procedure that segmentation methods apply to probability maps in order to create binary lesion masks. Currently, there are no stand-alone automated approaches for choosing thresholds for segmentation. After probability maps are created, experts may inspect each subject and visually determine a threshold to apply that performs well. Likewise, users may pick a single threshold that generally performs well across all subjects (Sweeney et al. 2013). These two thresholding methods, similar to manual segmentation, introduce human bias, cost, and time into the automated procedure. Several recent publications use cross-validation approaches for determining a threshold to apply to all subjects (see Roy et al. 2015; A. M. Valcarcel, Linn, Vandekar, et al. 2018 for example), but most methods do not provide sufficient detail to reproduce the thresholding approach. Further, these methods propose a group-level threshold rather than subject-specific thresholds.

In this paper we propose TAPAS, a Threshold Approach to Probability Map Automatic Segmentation, to address these and related problems. Using probability maps generated by an automatic segmentation method we fit the subject-specific threshold that yields maximum expected Sørensen’s-Dice Similarity Coefficient (*DSC*) based on a naive estimate of lesion volume using a general additive model. After training on a subset of subjects with manual segmentations, the TAPAS model can be applied to estimate a subject-specific threshold to apply to lesion probability maps in order to obtain automatic segmentations. This approach provides a generalizable method to subject-specific threshold detection by attempting to estimate a threshold that optimizes *DSC* and reduces bias. The TAPAS method is fully transparent, fast to implement, and simple to modify for new data sets.

## 2. Materials and methods

### 2.1 Data and preprocessing

The first data set studied (JHH data) was collected at the Johns Hopkins Hospital in Baltimore, Maryland. This data set consists of 98 subjects with MS; four were excluded due to poor image quality. Participants included were between 21.4 and 67.3 years of age, and 69 were women. Additionally, we had 1 subject diagnosed with clinically isolated syndrome, 9 subjects diagnosed with primary progressive MS, 60 subjects diagnosed with relapsing-remitting MS, and 24 subjects diagnosed with secondary progressive MS. Disease duration was defined as years since diagnosis. Additionally, subjects were examined by a neurologist to assess Expanded Disability Status Scale (EDSS) score. Patient demographics and disability scores are in **Table 1**; for more details see Sweeney et al. (2013). **Table 1** shows large variability in the manual T2 hyperintense lesion volumes.

**Table 1:** Demographic information for subjects in this study are provided. We included information from 94 patients imaged at Johns Hopkins’s Hospital (JHH) and 40 patients imaged at the Brigham and Women’s Hospital (BWH).

|  | Mean | Std. Dev. | Min. | Max. |
| --- | --- | --- | --- | --- |
| **JHH (n = 94)** |  |  |  |  |
| Age (years) | 43.4 | 12.3 | 21.4 | 67.3 |
| Disease duration (years) | 11.3 | 9.2 | 0.0 | 45.0 |
| Expanded Disability Status Scale score | 3.9 | 2.1 | 0.0 | 8.0 |
| Lesion volume (mL) | 11.5 | 13.1 | 0.0 | 77.0 |
| **BWH (n = 40)** |  |  |  |  |
| Age (years) | 50.4 | 9.9 | 30.4 | 69.9 |
| Disease duration (years) | 14.5 | 4.6 | 3.8 | 21.3 |
| Expanded Disability Status Scale score | 2.3 | 1.6 | 0.0 | 7.0 |
| Lesion volume (mL) | 13.6 | 12.8 | 0.6 | 52.0 |
| Timed 25-ft walk (seconds) | 11.5 | 6.9 | 1.0 | 25.0 |

For the JHH data, whole-brain 3D T1-weighted (T1), 2D T2-weighted fluid attenuated inversion recovery (FLAIR), T2-weighted (T2), and proton density-weighted (PD) images were acquired on a 3 Tesla (3T) MRI scanner (Philips Medical Systems, Best, The Netherlands). A more detailed description of the acquisition protocol was provided in previously published work (Sweeney et al. 2013; A. M. Valcarcel, Linn, Vandekar, et al. 2018). Manual T2 hyperintense lesion segmentations for each subject were delineated by an imaging scientist with more than 10 years of experience.

All images were N3 bias corrected (Sled, Zijdenbos, and Evans 1998), then the T1 scan for each subject was rigidly aligned to the Montreal Neurological Institute (MNI) standard template space at 1 *mm*^3^ isotropic resolution. FLAIR, PD, and T2 images were then aligned to the transformed T1 image. Extracerebral voxels were removed from all images using the Simple Paradigm for Extra-Cerebral Tissue Removal: Algorithm and Analysis (SPECTRE) algorithm (Carass et al. 2011). MRI scans were acquired in arbitrary units, and therefore analyzing images across subjects and imaging centers required that images be intensity-normalized. We thus intensity normalized each modality using *WhiteStripe* (Shinohara et al. 2014; Muschelli and Shinohara 2018). All image preprocessing was conducted using tools provided in Medical Image Processing Analysis and Visualization (MIPAV) (McAuliffe et al. 2001), TOADS-CRUISE (http://www.nitrc.org/projects/toads-cruise/), Java Image Science Toolkit (JIST) (Lucas et al. 2010), and R (version 3.5.0) (R Development Core Team 2018) software packages.

We used a second data resource collected at the Brigham and Women’s Hospital (BWH data) in Boston, Massachusetts from 40 subjects with MS. MRI data were consecutively obtained. Participants were between 30.4 and 69.9 years of age, and 28 were women. Additionally, we had 32 subjects diagnosed with relapsing-remitting MS and the remaining 8 subjects diagnosed with secondary progressive MS. Disease duration was defined as years since first symptoms. In order to assess the level of physical ability and ambulatory function, an MS neurologist examined patients to evaluate Expanded Disability Status Scale (EDSS) and timed 25-foot walk (T25FW) (in seconds). Patient demographics are provided in **Table 1** and further described in A. M. Valcarcel, Linn, Khalid, et al. (2018). Manual T2-hyperintense lesion volumes in this sample were less variable than the JHH data but still showed lesion load diversity.

For the BWH data, high-resolution 3D T1-weighted, T2-weighted, and fluid-attenuated inversion recovery (FLAIR) scans of the brain were collected on a Siemens 3T Skyra unit with a 20-channel head coil. The detailed scan parameters have been reported previously (Meier et al. 2018; A. M. Valcarcel, Linn, Khalid, et al. 2018). Two trained observers manually delineated T2 hyperintense lesions independently. After delineations were completed, the segmentations were then reviewed together in order to form a consensus. A senior experienced observer was consulted in the event of a disagreement. A single reviewer then manually segmented the T2 hyperintense lesions on the FLAIR image after all readers agreed on lesional presence in each voxel.

We performed N4 bias correction (Tustison et al. 2010) on all images and rigidly co-registered T1 and T2 images for each participant to the FLAIR at 1 *mm*^3^ resolution. Extracerebral voxels were removed from the registered T1 images using Multi-Atlas Skull Stripping (MASS) (Doshi et al. 2013) and the brain mask was applied to the FLAIR and T2 scans. We intensity-normalized images to facilitate across-subject modeling of intensities using *WhiteStripe* (Shinohara et al. 2014; Muschelli and Shinohara 2018). Image preprocessing was applied using software available in R (version 3.5.0) (R Development Core Team 2018) and from NITRC (https://www.nitrc.org/projects/cbica_mass/).

The Institutional Review Boards at the appropriate institutions approved these studies.

### 2.2 TAPAS algorithm

Although the two data sets were processed using different pipelines, the proposed technique is completely independent of the preprocessing pipeline. TAPAS simply relies on a continuous map of degree or probability of lesion at each voxel in the brain. Maps are generated by an automatic segmentation algorithm in order to predict a subject-level threshold for segmentation. In our experiments, we used the predicted lesion probability maps from a Method for Inter-Modal Segmentation Analysis (MIMoSA) (A. M. Valcarcel, Linn, Vandekar, et al. 2018; A. M. Valcarcel, Linn, Khalid, et al. 2018), an automatic segmentation procedure. We first divided the data set under study into two parts: the first is used for training TAPAS, and the second we refer to as the test set. In the training set of size *N/*2, we apply a grid of thresholds *τ*_1_, *…*, *τ*_*J*_, denoted as ***τ***, to the probability map in order to generate estimated lesion segmentation masks. For each subject in the training set we let ***τ*** vary from *τ*_1_ = 0% to *τ*_*J*_ = 100% by in 1% increments and calculate *DSC* between each estimated segmentation mask and the corresponding manual segmentation for the image. It is possible this step could be implemented using an optimization framework and may result in a reduction in computation time, but we did not validate other optimization approaches. Once lesion masks are generated after thresholding, we remove any lesions smaller than 8 *mm*^3^ (Shinohara et al. 2011; A. M. Valcarcel, Linn, Khalid, et al. 2018). We then estimated:

1. 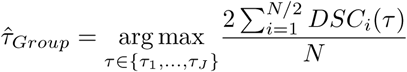, and
2. 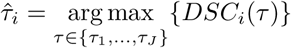 for each subject *i*.

The threshold estimated by 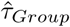 represents the threshold that produces maximum average *DSC* across all subjects in the training set, and 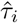 is defined as the subject-specific threshold that yields maximum *D SC* for s ubject *i*.

We apply 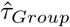 to each respective subject and obtain a naive estimate of the volume, 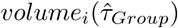. We then regress 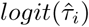 on the naive volume estimate, 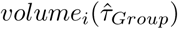, onusing a general additive model with a Gaussian link. The general additive model was chosen over linear models after manual inspection of scatter plots indicated non-linear trends. This is evident in the scatter plot displayed in the bottom left panel of **Figure 2**. We use a Gaussian link function since both 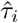 and 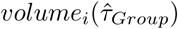 are continuous. Unfortunately, the Gaussian link does not bound the outcome 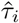 between 0 and 1; so, rather than modeling 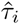, we model 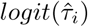 to force 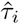 to be between 0 and 1. We implement the general additive model using the *gam* function available through the *mgcv* package in R. This function fits the model using a penalized scatter plot smoother with thin-plate splines and smoothing parameter estimated using generalized cross-validation (Wood 2003, n.d., 2004; Wood, Pya, and Säfken 2016). More specifically, the following general additive model is fit as the TAPAS model:

**Figure 2:**
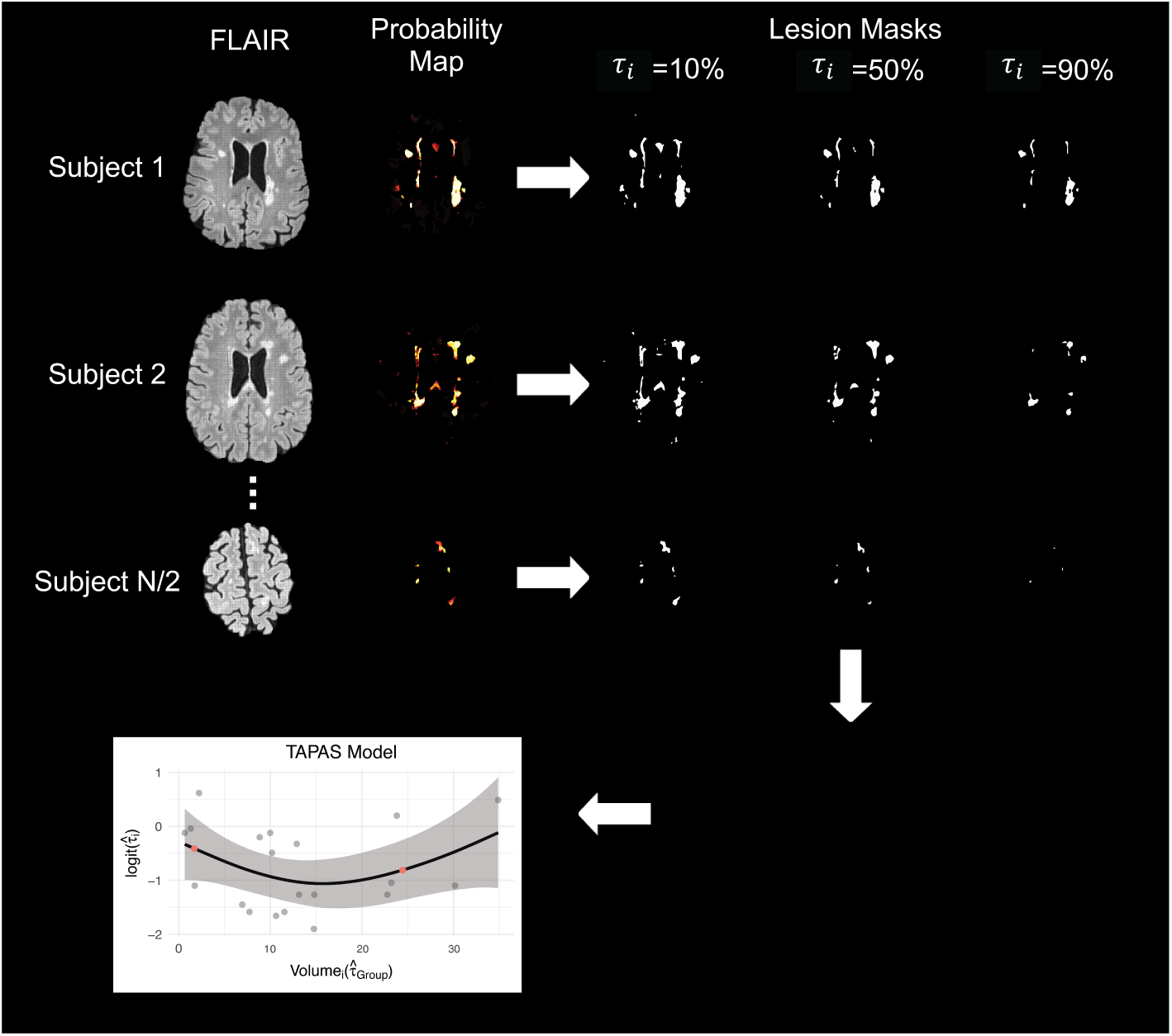
Axial slices of the TAPAS procedure are shown. A set of training scans with manual delineations were used to train and apply MIMoSA in order to obtain probability maps. For each subject’s probability map, we applied thresholds at *τ*_*i*_ = 0% to 100% by 1% to create estimated lesion masks. For simplicity, in this example, we have only shown *τ*_*i*_ = 10%, 50%, and 90%. Based on *DSC* calculations within and across subjects we calculated 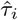 and 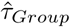 Using 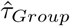 we obtained 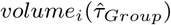 We fit the TAPAS model and applied it to subjects in the test set to determine 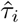. Red points in the plot represent 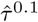 and 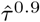, or lower and upper bounds at the volume associated with the 10th and 90th percentile.

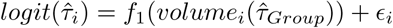

**where** *ϵ*_*i*_ ∼ *N* (0, ***σ***^**2**^).

In the model fitting procedure, we exclude subjects from model training if their 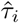 produces an estimated segmentation mask with *DSC <* 0.03. We found this to empirically improve TAPAS performance as it removes subjects for which even the best performing 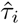 yields an inaccurate automatic segmentation mask.

After the TAPAS model is fit, we apply the model to subjects in the testing set. For each subject *i*, we obtain a probability map from an automatic segmentation procedure. We then use 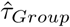 to threshold the probability map in order to estimate 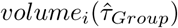. We use these predicted volumes in the TAPAS model to estimate the fitted value 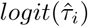, the subject-specific threshold. We re-threshold the probability maps by 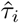 to generate the lesion segmentation mask. Similar to the training set, we also update these segmentation masks by removing any lesions smaller than 8 *mm*^3^ (Shinohara et al. 2011; A. M. Valcarcel, Linn, Khalid, et al. 2018). These updated masks are the final product of the TAPAS model, and can be used to obtain lesion metrics such as volume and count.

When applying the TAPAS model in the testing set, we aim to reduce extrapolation and excessive variability associated with left and right tail behavior of the spline model. Thus, for any volume we obtain using 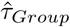 that is larger than the volume associated with the 90th percentile, we use the threshold for the subject whose volume is at the 90th percentile, denoted 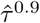, rather than the fitted 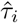 Similarly, for any volume we obtain from 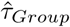 that is smaller than the volume associated with the 10th percentile, we use the value of 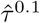 **Figure 2** shows an outline of the full TAPAS procedure and model.

To implement TAPAS, we developed an R package that is available with documentation on GitHub (www.github.com/avalcarcel9/tapas) and Neuroconductor (https://neuroconductor.org/package/rtapas).

### 2.3 Performance assessment

For the two data sets in this study (JHH and BWH), we ran separate Monte Carlo-resampled split-sample cross-validations. More specifically, we repeatedly sampled (100 times) without replacement to assign half of the subjects in the study to each of the training and testing sets. In each training set, we applied MIMoSA using the R package *mimosa* (A. Valcarcel 2018) available on Neuroconductor (https://neuroconductor.org/package/mimosa) (Muschelli et al. n.d.). After fitting the MIMoSA model using subjects in the training set, we generated probability maps for all subjects in the training and testing sets.

In each split-sample experiment, the training set was used to fit the TAPAS model and the testing set applied the TAPAS model to determine a subject-specific threshold 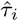. This subject-specific threshold was used to create binary lesion segmentation masks and calculate lesion volume. We compared the TAPAS-generated masks and volumes, denoted by the subscript *TAPAS*, whereas masks and volumes generated by the 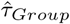 threshold are henceforth denoted with the subscript *Group*. The use of 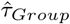 to threshold probability maps and generate lesion segmentations was previously applied (A. M. Valcarcel, Linn, Vandekar, et al. 2018; A. M. Valcarcel, Linn, Khalid, et al. 2018) and aided in automatic segmentation measures compared to user-defined threshold application.

We provide quantitative comparisons between TAPAS and the group thresholding procedure for subjects in the testing set. First, we compared the correlation between *volume*_*TAPAS*_ or *volume*_*Group*_ and *volume*_*Manual*_ within each split-sample experiment and averaged across folds to assess the correspondence between volumes. We denote the correlation estimates by 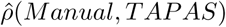 and 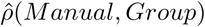. Second, to assess whether segmentation masks produced using TAPAS or the group thresholding procedure differed in accuracy as measured by *DSC*, we compared segmentations between lesion masks produced by TAPAS (*DSC*_*TAPAS*_) and those produced by the group thresholding procedure (*DSC*_*Group*_) with manual segmentations. We compared these measures using a paired t-test within each split-sample experiment using subjects in the test set. Third, to assess bias and inaccuracy present in *volume*_*TAPAS*_ and *volume*_*Group*_ we calculated absolute error defined as *AE* = |*Threshold V olume - M anual V olume|*. In order to determine whether *AE* differed statistically, paired t-tests were conducted between *AE*_*TAPAS*_ and *AE*_*Group*_ within each split-sample experiment.

To adjudicate whether TAPAS yielded volumetrics with similar phenotype associations, we calculated the Spearman’s correlation coefficient between *volume*_*TAPAS*_, *volume*_*Group*_, and *volume*_*Manual*_ and clinical variables. We denote these correlations by 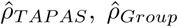, and 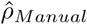 respectively. We estimated correlations in each split-sample experiment and averaged across experiments.

## 3. Results

### 3.1. Volumetric bias assessment

Using Bland-Altman visualization, we compare automatic and manual volumes in **Figure 3**. Subject-level volumes were obtained by averaging each subject’s measurement for all split-sample experiments in which it was allocated to the testing set. The JHH data *volume*_*Group*_ estimate exhibits systematic bias, evident in **Figure 3** for volumes exceeding 20 mL. Visually, we observed a moderate inverse relationship in these subjects. This indicates that *volume*_*Group*_ underestimated *volume*_*Manual*_ in subjects with larger lesion loads with increasing magnitude. Unlike the Group Bland-Altman plot, the TAPAS plot does not exhibit obvious patterns of systematic bias. The cluster of points that begins to negatively deviate from 0 in the Group plot are still centered randomly around 0 in the TAPAS plot. Additionally, the mean and standard deviation for the differences is smaller using *volume*_*TAPAS*_ compared to *volume*_*Group*_. There are four points that lie outside the limits of agreement in both thresholding procedures, but, in the TAPAS plot, these are closer to 0.

**Figure 3:**
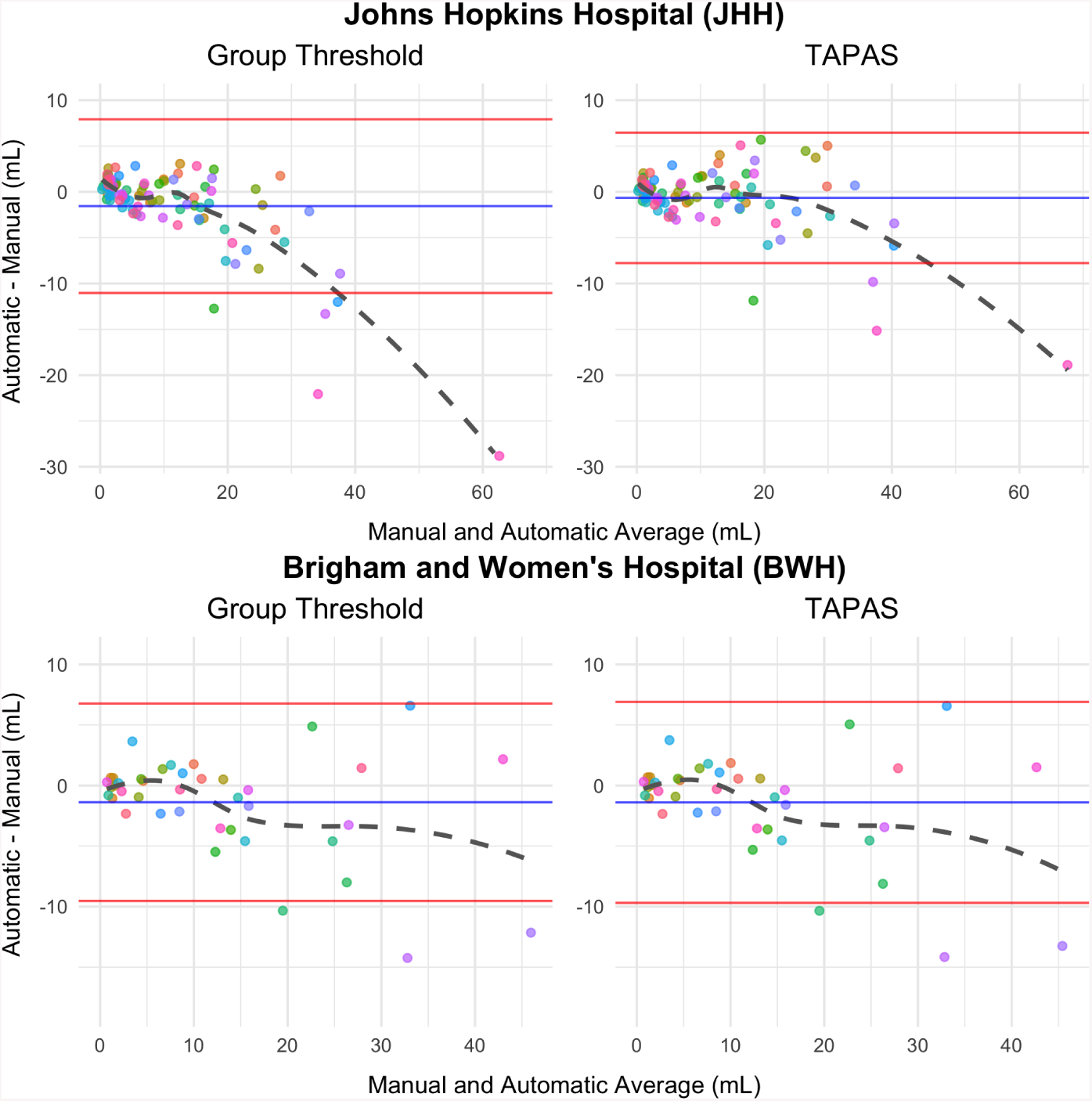
Bland-Altman plots comparing *volume*_*Manual*_ with automatic thresholding approaches (*volume*_*Group*_ or *volume*_*TAPAS*_) are shown. The mean of the difference in volume is presented in blue and the mean plus and minus the standard error is shown in red. Each point represents a unique subject. Subject-specific points were obtained by averaging results across test set subjects in each split-sample fold.

The BWH Bland-Altman plots are nearly identical and almost indistinguishable when comparing the group threshold procedure with the TAPAS outputs. There does not appear to be a systematic bias in either *volume*_*Group*_ or *volume*_*TAPAS*_ estimates since points are randomly scattered around 0 in the positive and negative directions. This exemplifies TAPAS’s propensity to conserve unbiased estimates when systematic bias is absent.

### 3.2 Absolute error assessment

Scatter plots and predicted linear models are presented in **Figure 4** to compare the absolute error (*AE*_*TAPAS*_ and *AE*_*Group*_) between *volume*_*Manual*_ and *volume*_*Group*_, respectively. The JHH data plot showed smaller absolute error estimates associated with *volume*_*TAPAS*_ compared to *volume*_*Group*_. This is highlighted by the negative shift in *AE*_*TAPAS*_ points throughout as well as smaller slope estimates provided in the top left corner. The coefficient associated with *AE*_*Group*_ is 0.27 while the coefficient associated with *AE*_*TAPAS*_ is 0.18. Using these coefficients, for a unit increase in *volume*_*Manual*_, *AE*_*TAPAS*_ is predicted to be 0.09 mL less than *AE*_*Group*_. In the BWH data, all values were remarkably similar across the methods. The results in **Figure 3** and **Figure 4** are consistent and indicate that *volume*_*TAPAS*_ is less biased than *volume*_*Group*_.

**Figure 4:**
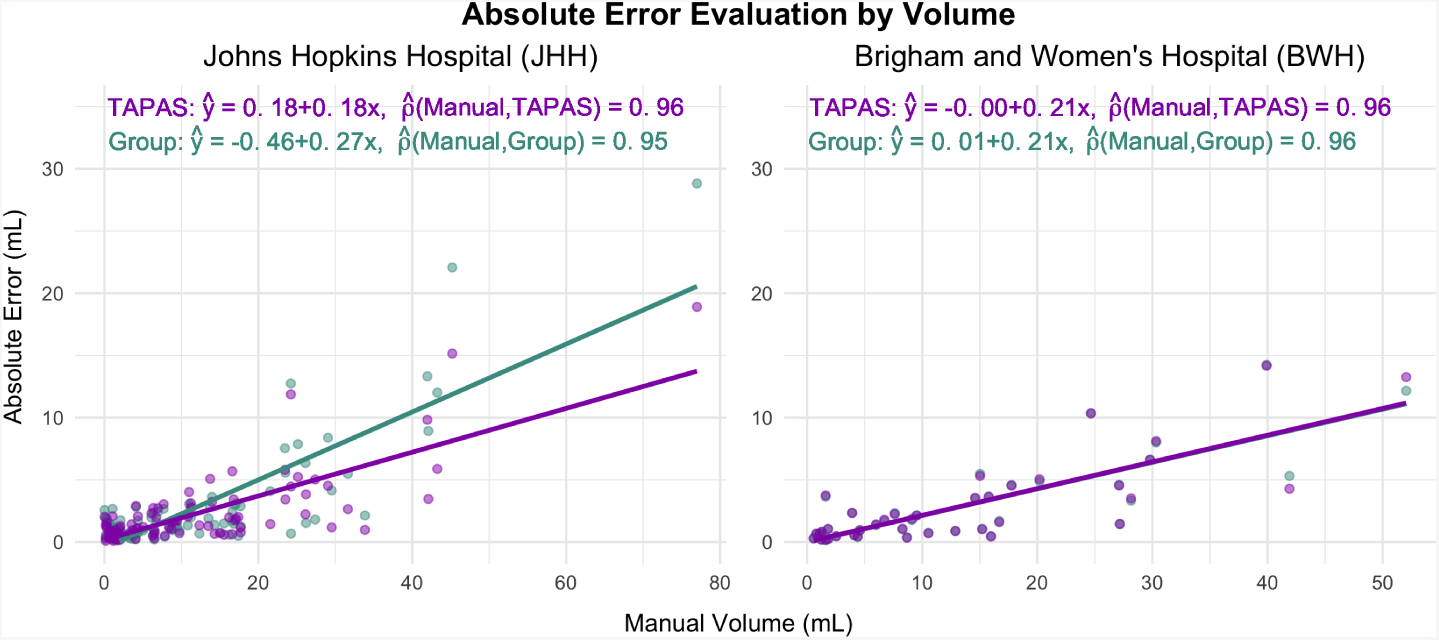
Scatter plots with fitted linear models are presented for the subject-level average absolute error (*ŷ*) on manual volume (x) in mL. Fitted equations are given in the top left corner. Additionally, the correlation estimates between manual volume and respective threshold produced volume are provided.

The average *AE* across subjects in the testing sets and iterations in the JHH data is 2.21 mL using the TAPAS subject-specific threshold compared to 2.7 mL using the group thresholding procedure. In the BWH data, the average *AE* using TAPAS is 2.87, while using the group thresholding procedure generates an average *AE* of 2.86. TAPAS yields equal or reduced average *AE*. The average *DSC* across subjects in the testing sets and iterations in the JHH data is 0.61 using the TAPAS subject-specific threshold compared to 0.6 using the group thresholding procedure. In the BWH data, the average *DSC* is 0.66 for both TAPAS and the group thresholding procedure. TAPAS yields equal or superior average *DSC*.

To examine this statistically, we employed one-sided paired t-tests to evaluate *AE* and *DSC* from TAPAS compared with those obtained from the group thresholding procedure. **Figure 5** shows violin plots of p-values from both sets of tests for the two data sets. The labels beneath each violin show the number of p-values less than *α* = 0.05 that favor the TAPAS measure (i.e. a reduction in *AE* and an increase in *DSC*). In the JHH data, there was a skew towards smaller p-values. More than half of the split-sample experiments resulted in p-values below the *α* = 0.05 for *AE* and *DSC*. This indicates superior performance using TAPAS compared to the group thresholding procedure. The BWH data was more uniform with approximately a tenth of p-values favoring TAPAS. P-values above the *α* = 0.05 threshold only inferred no difference in TAPAS and group thresholding measures.

**Figure 5:**
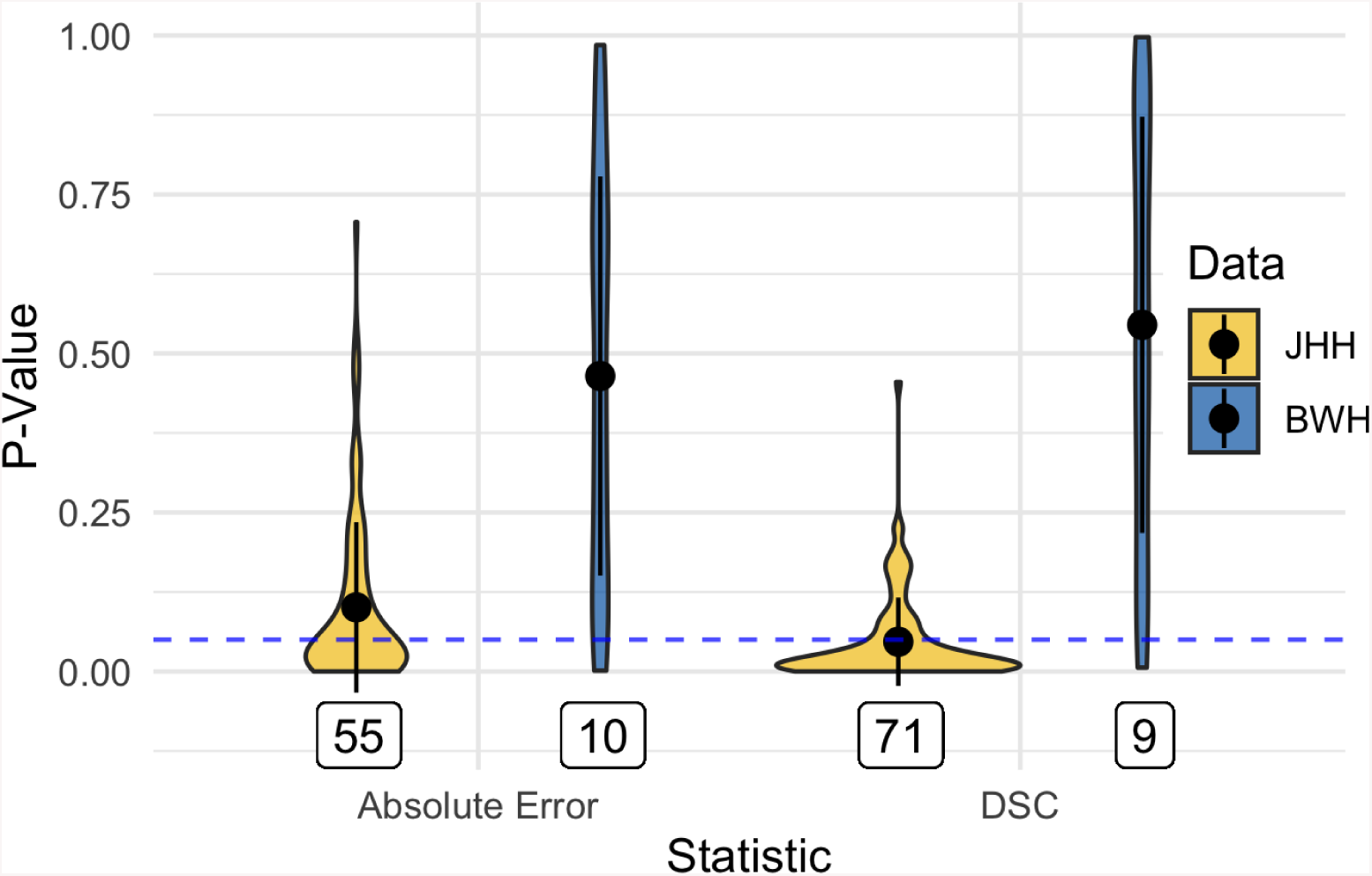
Violin plots of p-values from paired t-tests to compare subject-level absolute error (*AE*) and Sørensen-Dice coefficient (*DSC*) in each test set are presented. The mean for each statistic and data set is presented as points within each violin plot and the black lines extend the mean by the standard deviation. Labels below represent the number of significant p-values favoring TAPAS performance measures. The dashed horizontal blue line highlights the *α* = 0.05 cutoff.

To be thorough and transparent, we counted the number of split-sample folds in which t-tests concluded in favor of using the group threshold. The JHH data did not result in any iterations concluding group threshold superiority. In the BWH data, group threshold tests favored the TAPAS procedure in approximately a tenth of folds.

### 3.3 Correlation analysis

Comparisons between volume estimates are provided in the top left corner of **Figure 4** to the right of the fitted linear models. It is important to note that this correlation estimate is not related to the fitted linear models but rather Spearman’s correlation between *volume*_*Manual*_ and *volume*_*TAPAS*_ or *volume*_*Group*_. Interestingly, 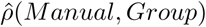 and 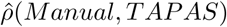 are nearly identical with 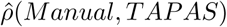 only slightly higher.

In addition to the volumetric correlation analyses, we assessed the relationship between *volume*_*TAPAS*_, *volume*_*Group*_, and *volume*_*Manual*_ with various clinical variables. These results are provided in **Table 2**. All correlations found are modest but aligned with previously published literature (A. M. Valcarcel, Linn, Khalid, et al. 2018; Stankiewicz et al. 2011; Barkhof 1999; Tauhid et al. 2014). In the JHH data 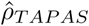 and 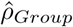 are indistinguishable from each other and slightly larger than 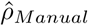. Similarly, the BWH data show identical 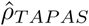 and 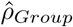 nearly equivalent to 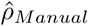. In terms of phenotypic associations *volume*_*TAPAS*_ yielded similar correlation estimates as *volume*_*Group*_ and *volume*_*Manual*_.

**Table 2:** Subject-specific volume estimates, *volume*_*Manual*_ (Manual), *volume*_*TAPAS*_ (TAPAS), and *volume*_*Group*_ (Group), were compared with clinical covariates available in each data set and are represented in this table. Spearman’s correlation coefficient was obtained in the testing set for each iteration and averaged across folds. Clinical variables included Expanded Disability Status Scale (EDSS) score, disease duration in years, and timed 25-ft walk (T25FW).

| | Estimates for $\hat{\rho}$ | | |
| --- | --- | --- | --- |
|  | Group | TAPAS | Manual |
| **JHH** |  |  |  |
| EDSS | 0.36 | 0.36 | 0.30 |
| Disease Duration | 0.41 | 0.40 | 0.40 |
| **BWH** |  |  |  |
| EDSS | 0.40 | 0.40 | 0.43 |
| Disease Duration | 0.30 | 0.30 | 0.27 |
| T25FW | 0.02 | 0.02 | 0.03 |

### 3.4 Threshold evaluation

In **Figure 6** there are a few notable differences between the threshold scatter plots produced from TAPAS and those produced by the group thresholding procedure. In both data sets the subject-specific thresholds have a much wider range than the group thresholds. In the JHH data, the distribution shape is bimodal for the subject-specific thresholds but uni-modal for the group thresholds. In the BWH data, the distribution shape is similar between the two thresholding approaches.

**Figure 6:**
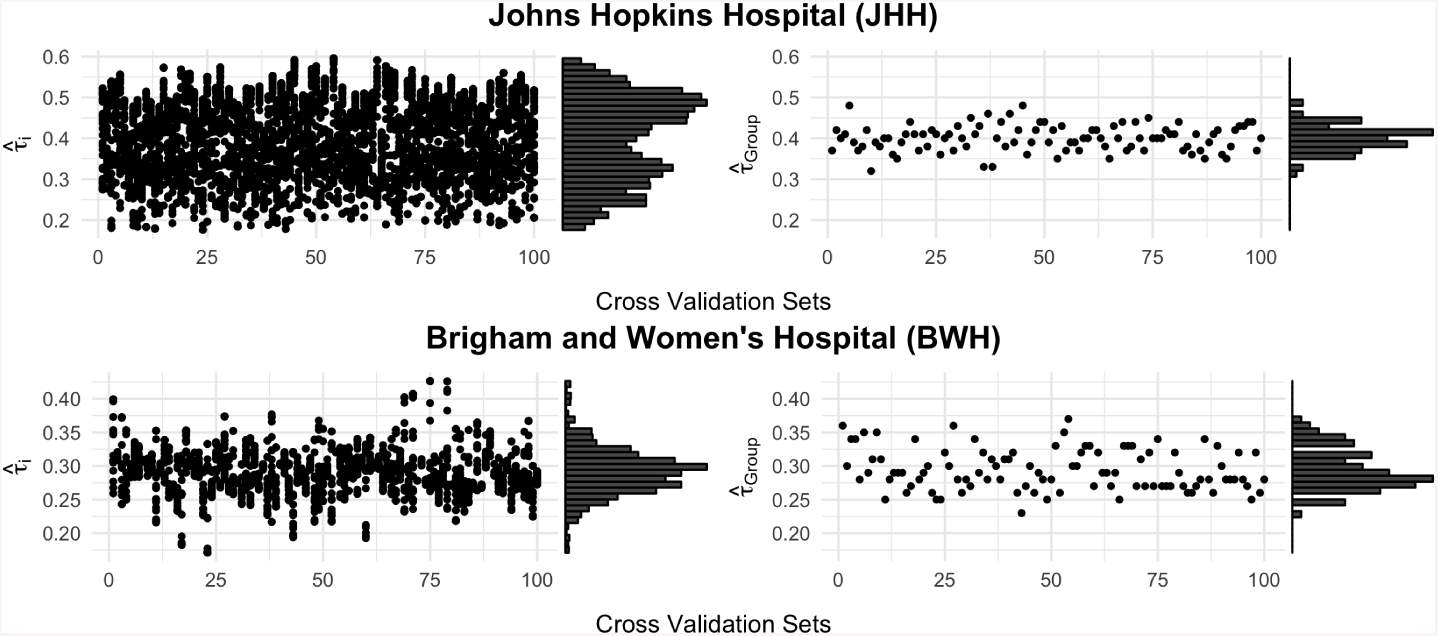
Scatter plots of the subject-specific threshold 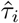 and 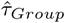 (group thresholding procedure) on cross-validation number are presented with marginal histograms for both data sets.

We also plotted the residuals for 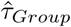 and 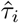 separately against manual volume in **Figure 7**. The residual is defined as the difference between the true subject-specific threshold that maximizes *DSC* with the manual segmentation for subject *i, τ*_*i*_, and the threshold determined from either TAPAS or the group thresholding procedure. The JHH data TAPAS residual plot shows no obvious pattern indicating the model fit well. The group thresholding procedure displays a general decreasing pattern. For subjects with small lesion loads, 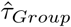 underestimates *τ*_*i*_, while for subjects with larger lesion loads, 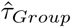 overestimates *τ*_*i*_. The clear pattern indicates systematic bias in the threshold approach and generally worse individual level predicted thresholds. The BWH data plots are essentially identical with very subtle differences. In these data, both thresholding approaches appear to fit the data well since points are randomly dispersed around 0 with no notable pattern. Using the residual ranges on the y-axes in both data sets, we see a wide spread of residuals. These values also give insight into the diversity of predicted thresholds from both thresholding approaches.

**Figure 7:**
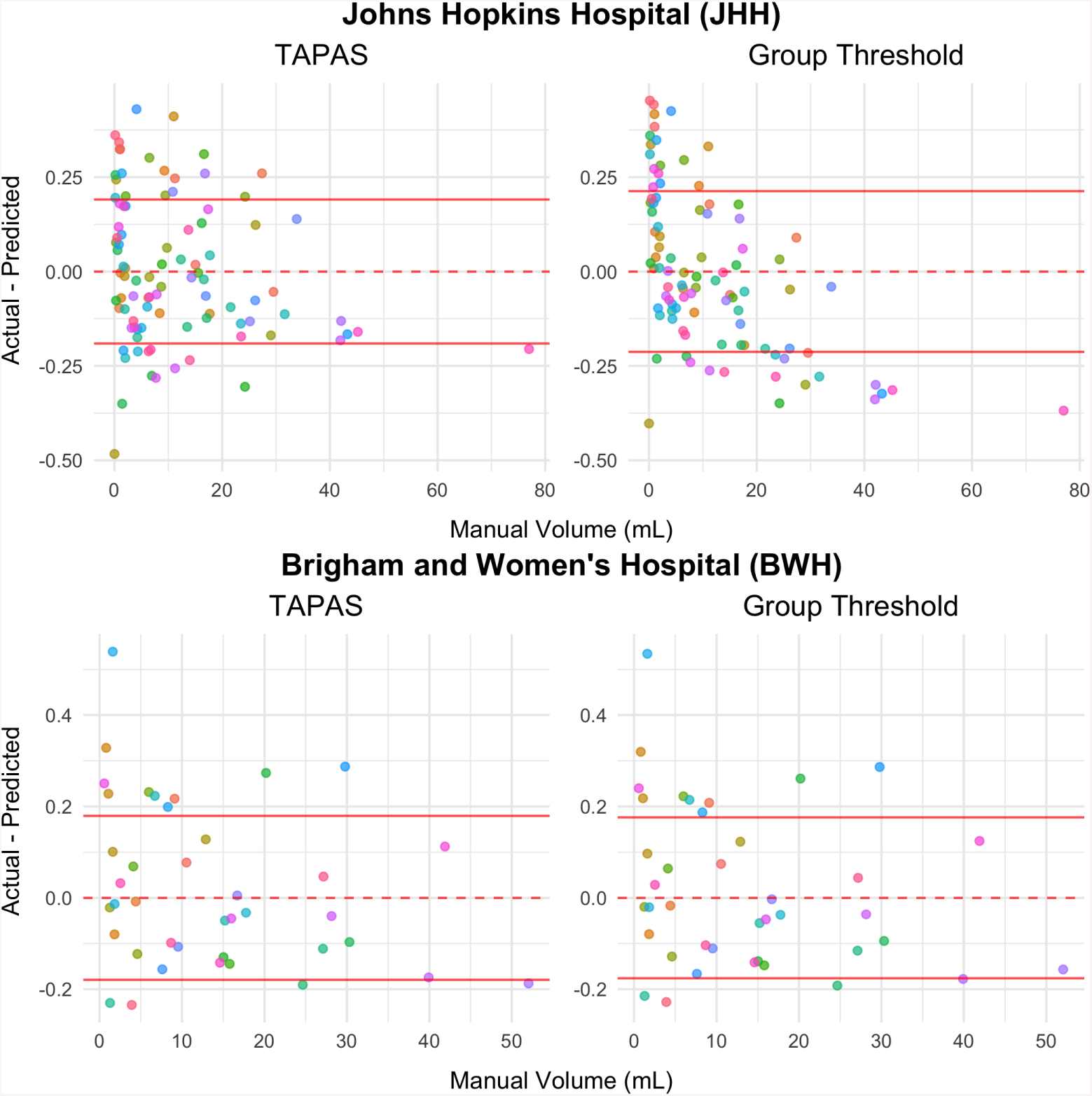
Plots depicting threshold residuals against manual volume are presented. The residual is defined as the difference between the actual subject-specific threshold that maximizes subject-level *DSC, τ*_*i*_, and 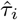 or 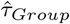. The dashed line highlights *y* = 0 while solid red lines represent *y* = 0 plus and minus the residual standard deviation.

### 3.5 Qualitative results

We present segmentations from the TAPAS and group thresholding approaches as well as manual delineations in **Figure 8**. This figure shows that TAPAS and the group thresholding procedure generally agree with the manual segmentation. Some tissue was manually segmented and not detected by either thresholding algorithm. The major differences between all the methods are found at the boundaries of lesions, which are known to be difficult to discern for both automatic and manual approaches. Overall, the automatic segmentation algorithm paired with either thresholding approach is able to detect majority of lesional space with few false positive area.

**Figure 8:**
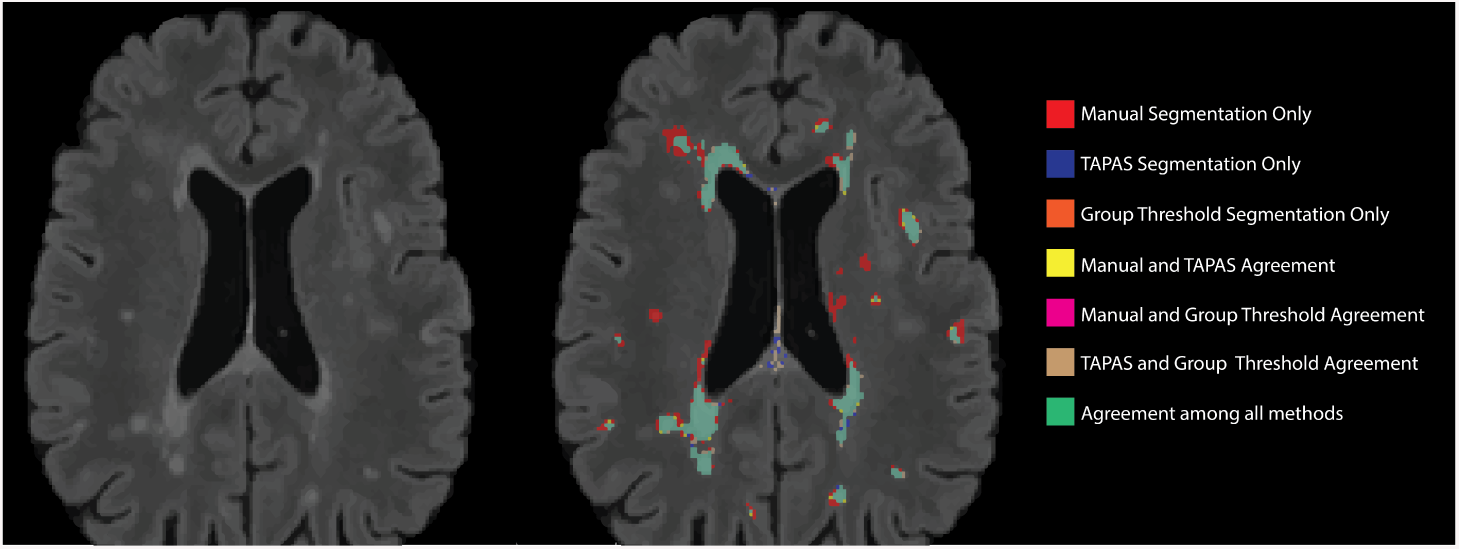
T2 hyperintense lesion segmentations from an example axial slice are displayed. The colors represent the different individual or overlapping segmentations obtained from manual, TAPAS threshold, and group threshold masks. The majority of segmented area was in agreement among all lesion masks (green). Both the group thresholding approach and TAPAS missed some area that was manually segmented (red). There was a small amount of area where TAPAS and manual segmentations agreed (yellow), but almost no area where only the group threshold agreed with the manual segmentation (fuchsia).

## 4. Discussion

Most automatic segmentation algorithms produce continuous maps of lesion likelihood, which are subsequently thresholded to create binary lesion segmentation masks. While a number of automatic approaches exist for lesion segmentation, there are few automatic algorithms available for threshold detection. Thresholds are commonly chosen using cross-validation procedures conducted at the group level, or arbitrarily through subjective human input. This introduces variability and biases in automatic segmentation results. Furthermore, thresholding approaches often apply a single common threshold value to all subjects’ probability maps. This lack of subject specificity may lead to inaccuracy in lesion segmentation masks, especially in subjects with the smallest and largest lesion loads.

This study sought to address these issues by introducing a fully automated algorithm for subject-specific threshold prediction that also reduces volumetric bias if present. The TAPAS procedure is easily implemented and performs well on data acquired with different scanning protocols or pre-processed with different pipelines. We validated TAPAS in two unique data sets from different imaging centers using 3T MRI scanners from different vendors.

The TAPAS procedure is a fully automated thresholding approach that determines a subject-specific threshold to apply to continuous maps (including predicted probability maps) for automatic lesion segmentation. TAPAS volume estimates are accurate and reduce systematic biases associated with differential total lesion load when present. In the JHH data, we observed such a bias using the MIMoSA algorithm, which was mitigated using TAPAS.

The BWH data used a consensus approach with two trained raters to manually segment lesions. We believe this approach reduces intra- and inter-rater variability normally present with a single rater and allows for a closer approximation of the ground truth, and, thus, better training of automatic approaches. The Bland-Altman plots in these data indicate unbiased estimation using a group threshold or TAPAS. In this study without systematic volumetric biases, we showed that TAPAS preserves the unbiased volumetric estimation of the automated segmentation technique.

In clinical trial evaluations of therapeutic effectiveness, associations between clinical variables and lesion volume are of primary interest. TAPAS and group threshold volumes resulted in similar correlations to clinical variables as the manual volume. This supports the validity of the proposed automatic segmentation and thresholding procedures.

TAPAS is a post-hoc subject-specific threshold detection algorithm built to reduce volumetric bias associated with automatic segmentation procedures. In this study, we optimized TAPAS using *DSC* though other measures are possible if validated. For example, absolute error or mean square error may be more meaningful in other settings. In fact, we explored minimizing absolute error in early explorations but found *DSC* to slightly outperform absolute error. Automatic approaches are constantly being built and improved upon to yield more accurate and robust methods. TAPAS allows for improvement upon even the most accurate and robust automatic segmentation procedures with no observed addition of error. Beyond MS or MRI, this methodology can be used for automatic segmentation of other tissues or body parts using different imaging types after proper validation.

There are several notable limitations to the proposed algorithm. First, the method must be used in conjunction with continuous maps of likelihood of lesion, so investigators must use automatic approaches that generate these maps for adaptive thresholding. Further, in this work we evaluated the TAPAS procedure with only a single automatic segmentation approach, MIMoSA, applied to two data sets. Second, since the TAPAS model fits a generalized additive model, training data sets with small sample size, uniform lesion load, or those dissimilar from testing data may have a poor model fit or inappropriate threshold estimation. Furthermore, to apply TAPAS to longitudinally acquired data, such as those presented in the 2015 segmentation challenge, a sufficiently large sample of subjects with variable lesional volume is required.

Future developments will include specialized methods for the analysis of longitudinal lesion volumetrics. Additionally, we will validate TAPAS using other automatic segmentation approaches for MS lesion detection. The distribution of probability maps using other automatic approaches may differ and gains using TAPAS are unknown. In the implementation of TAPAS with other automatic segmentation approaches investigators should cross-validate the TAPAS procedure to ensure no losses in segmentation performance. It is possible that the underlying method may benefit from dynamic thresholds for smaller lesions and larger lesions even within the same subject. That is, we may need to move beyond even a subject-specific threshold since, when a subject has larger lesions, the error associated with larger lesions contributes more to the DSC metric than the same relative error associated with smaller lesions. There may thus be a tendency of TAPAS to better segment larger lesions at the cost of doing worse on smaller lesions.

## 5. Acknowledgements

The authors would like to thank Ciprian Crainiceanu for providing useful feedback concerning the model development. This work was supported by the National Institutes of Health R01NS085211, R21NS093349, R01MH112847, R01NS060910, R01EB017255, R01NS082347, R01EB012547, 2R01NS060910-09A1, NIND 2037033; and the National Multiple Sclerosis Society, RG-1507-05243, RG-1707-28586. The content is solely the responsibility of the authors and does not necessarily represent the official views of the funding agencies.

